# The impact of experimental design choices on parameter inference for models of growing cell colonies

**DOI:** 10.1101/171710

**Authors:** Andrew Parker, Matthew J. Simpson, Ruth E. Bakera

## Abstract

To better understand development, repair and disease progression it is useful to quantify the behaviour of proliferative and motile cell populations as they grow and expand to fill their local environment. Inferring parameters associated with mechanistic models of cell colony growth using quantitative data collected from carefully designed experiments provides a natural means to elucidate the relative contributions of various processes to the growth of the colony. In this work we explore how experimental design impacts our ability to infer parameters for simple models of the growth of proliferative and motile cell populations. We adopt a Bayesian approach, which allows us to characterise the uncertainty associated with estimates of the model parameters. Our results suggest that experimental designs that incorporate initial spatial heterogeneities in cell positions facilitate parameter inference without the requirement of cell tracking, whilst designs that involve uniform initial placement of cells require cell tracking for accurate parameter inference. As cell tracking is an experimental bottleneck in many studies of this type, our recommendations for experimental design provide for significant potential time and cost savings in the analysis of cell colony growth.

## 1. Introduction

The study of how cell populations grow and spread is integral to understanding and predicting the invasion of cancer, the speed of wound repair and the robustness of embryonic development [9, 24, 28]. However, the extent to which cell populations grow and spread is governed by multiple processes, including motility, proliferation, adhesion and cell death, making it difficult to elucidate the relative contributions of these processes to the growth and invasion of a cell colony [31]. As such, *in vitro* cell biology assays are routinely used to probe the mechanisms by which cells interact, and the key processes involved in the growth and expansion of cell colonies. These *in vitro* assays generally involve seeding a population of cells on a two-dimensional substrate, and observing the population as the individual cells move and proliferate and the density of the monolayer increases towards confluence. A useful approach to interpret the results of these assays involves using a mathematical model that incorporates mechanistic descriptions of processes such as cell motility and proliferation. By parameterising and validating the models using quantitative data from *in vitro* assays it is possible to provide quantitative insights into the mechanisms driving the growth and spreading of a cell population, and make experimentally testable predictions. However, it is not always clear how best to choose the experimental design, nor which summary statistics of the data to collect, in order to accurately and efficiently parameterise and validate models.

In this work we utilise a two-dimensional lattice-based exclusion process model that incorporates both motility and proliferation mechanisms. Our goal is to assess how our ability to accurately infer model parameters is affected by changes in the experimental design. Parameter inference is performed in a Bayesian framework using approximate Bayesian computation (ABC), allowing us to quantify the uncertainty of our parameter estimates and bypass the need to compute a likelihood function for the mechanistic model. By quantifying the information gain using the different experimental protocols, we are able to provide guidelines for experimental design in terms of the selection of experimental geometry and the collection of relevant quantitative summary statistics from imaging data.

### 1.1. Experimental design

Typically, there are two main types of two-dimensional *in vitro* experiments that are considered at the level of the population. The first experiment, shown in Figure 1(a), is often referred to as a *growth*-*to*-*confluence assay*. Here we observe a population of cells seeded, initially at low density, as the cells move and proliferate and the population increases in number to eventually occupy the whole domain under observation [5, 8]. The second experiment we consider is shown in Figure 1(b), and is often referred to as a *scratch assay*. It involves perturbing a population of cells at, or near, confluence by scraping away a region of the population and observing the resulting spread of the population into this, now empty, region [18, 33]. In this work we will explore the extent to which models of these two types of assays can be parameterised using various summary statistics of these experimental data.

**Figure 1:**
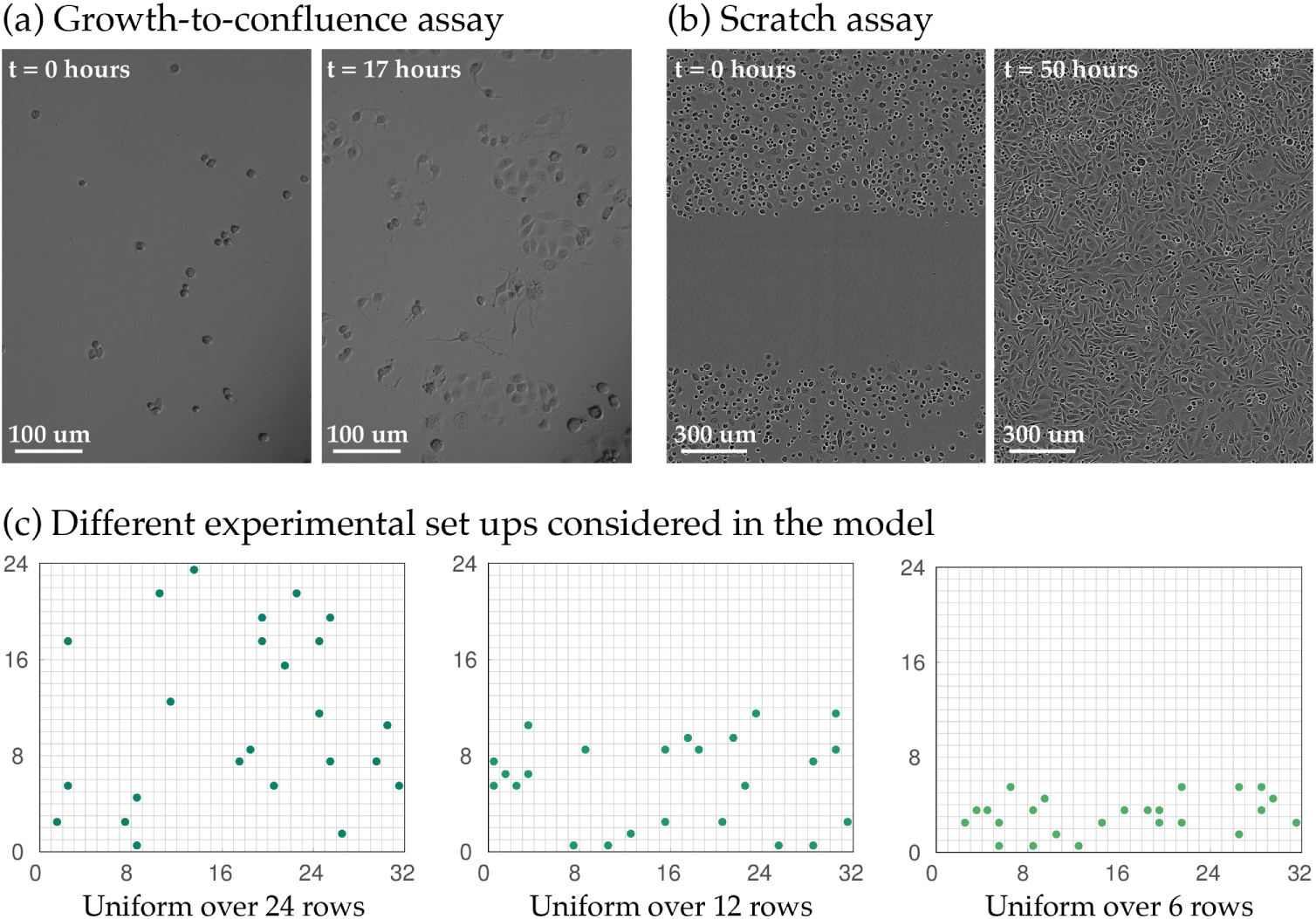
Two experimental designs considered in this work. (a) Images from a growth-to-confluence using MDA MB 231 breast cancer cells. See [25] for more information. (b) Images from a scratch assay with PC3 prostate cancer cells. See [11] for more information. (c) Schematics of the model. We use an on-lattice model, as described in the text, and vary the initial condition to replicate the experiments in (a),(b): we seed cells uniformly at random over 24 rows (left), 12 rows (middle) or six rows (right). The images in (b) are reproduced from Simpson et al. [25] with kind permission, whereas the images in (c) are reproduced from Johnston et al. [11] with kind permission.

### 1.2. Approximate Bayesian computation and summary statistics

Parameter inference is approached generally in one of two ways, either through a frequentist approach or a Bayesian approach [2, 6]. In frequentist inference, one generally seeks a point estimate of a parameter through maximum likelihood estimation, and captures uncertainty in the estimate through the generation of confidence intervals. A Bayesian approach instead derives a predictive posterior distribution for the model parameters ***θ*** given observed data 𝓓^*obs*^ [10]. The posterior, ℙ(***θ***|𝓓^*obs*^), satisfies ℙ(***θ***|𝓓^*obs*^) ∝ 𝓛(𝓓^*obs*^|***θ***) *π*(***θ***), where 𝓛(𝓓^*obs*^|***θ***) is the likelihood of obtaining the data from the given model and parameters, and the prior, *π*(***θ***), captures any previous knowledge of the parameters. For the mechanistic model we use in this work, the likelihood is intractable and so we use ABC to generate an approximate posterior distribution.

ABC has been used previously in the biological sciences, particularly to target complex problems in systems biology [13, 16], population genetics [4], and ecology [29]. There is also growing popularity in the use of ABC specifically in the investigation of cell biology processes [12, 22, 32]. ABC relies on repeated simulation of the model using parameters from the prior distribution, and the acceptance of these parameters sets whenever the model output is sufficiently close to the experimental data. These accepted parameter sets are then used to estimate the posterior distribution. In using ABC in practice there are several important user choices, most importantly the means of comparison between simulated and experimental data, typically through summary statistics and a suitable distance metric, and the threshold used for accepting or rejecting parameter sets [27, 30]. In particular, the summary statistics represent lower dimensional descriptions of the data and their choice is vitally important as they impact the amount of information gained about model parameters through the use of ABC [3].

The major new insights provided in this work are a quantitative understanding of how both the choices of experimental geometry and summary statistics impact the quality of the posteriors generated using ABC. The use of ABC rejection allows us to compare the quality of the posteriors resulting from a wide range of possible summary statistics in a computationally efficient way, and we quantify the information gain using the Kullback-Leibler divergence [14]. We also demonstrate how data-cloning ABC (ABC-DC) [21] can potentially be used to obtain maximum likelihood estimates (MLEs) of model parameters more efficiently than ABC rejection. Instead of targeting the likelihood of the observed data, the data cloning approach targets the likelihood corresponding to a large number of copies (also known as clones) of the data, where each data clone is assumed independent of the others. Data cloning results in a posterior distribution that has the MLE as its mean, and the variance can be related to the asymptotic variance of the MLE [15]. ABC-DC uses ABC Markov chain Monte Carlo with data cloning to facilitate convergence towards a MLE [21].

### 1.3. Aims and outline

The aim of this work is to demonstrate the use of Bayesian inference methods to characterise the motile and proliferative behaviour of individual cells within growing cell colonies. In particular, we aim to estimate parameters of a lattice-based volume exclusion model for cell colony growth in a variety of experimental geometries. In addition, we demonstrate the use of ABC-DC for more efficient maximum likelihood estimation. In Section 2 we introduce our model, and the various ABC algorithms and summary statistics we employ in this work. In Section 3 we assess how experimental design choices impact the quality of estimated posterior distributions, using both ABC rejection and ABC-DC, and we conclude with a discussion of our results in Section 4.

## 2. Methods

### 2.1. Mechanistic model

We employ a simple two-dimensional lattice-based exclusion process model akin to that of Simpson *et al*. [26], whereby *N*(*t*) cells occupy a square lattice with *R* rows and *C* columns at time *t*. During each time step of duration *τ*, we choose *N*(*t*) cells at random with replacement to attempt a movement or proliferation event into orthogonally adjacent lattice sites with probabilities *P_m_* and *P_p_*, respectively. For each cell we draw a uniform random number, *r* ~ 𝓤(0,1). If *r* ≤ *P_m_* the cell attempts a movement event into one of the four orthogonally adjacent lattice sites with equal probability, and if *P_m_* < *r* ≤ *P_m_* + *P_p_* the cell attempts a proliferation event, whereby a daughter cell is placed into one of these lattice sites, each with equal probability. If *P_m_* + *P_p_* < *r* ≤ 1 no movement or proliferation event is attempted. If a cell attempts to move or to place a daughter cell into an occupied lattice site, or outside of the domain, the attempted movement or proliferation event is aborted. To replicate experimental conditions, we take *R* = 24, *C* = 32, where lattice sites have length Δ = 18.75 *μ*m (corresponding to the approximate cell diameter of the cells considered in typical experiments). Simulations are initialised with cell positions randomly distributed in the first *R̂* rows of the domain, where *R̂* is chosen to mimic potential experimental conditions. To interpolate between the growth-to-confluence and scratch assay designs, we choose three initial conditions (see Figure 1(c)) initialising cells uniformly at random across either 24, 12 or six rows of the domain.

### 2.2. In silico data

Since our aim in this work is to better understand how experimental design impacts upon our ability to infer model parameters, we use our mechanistic model to generate *in silico* (observed) data that closely replicates that available from experiments (see Figure 1(a),(b)) and attempt to infer the parameters used to generate these *in silico* data. We use a time step of *τ* = 1/24 hr, and model parameters *P_m_* = 0.25 and *P_p_* = 0.0025 (motivated by estimates for the diffusivity and proliferation rates from similar experiments [11]). We record the final positions of cells in our *in silico* experiments after 12 hours, equating to 288 time steps. We also record trajectory data for five randomly chosen cells by recording their positions every eight time steps (corresponding to every 20 minutes [25]). As experiments are typically repeated several times to ensure reproducibility of results [25], we repeat simulations *M* = 10 times and average the resulting statistics.

### 2.3. Inference

We use ABC to estimate posterior distributions for the model parameters, ***θ*** = (*P_m_*,*P_p_*). ABC rejection is performed by repeatedly sampling from the prior distribution, *π*(***θ***), simulating the model, and accepting parameters that result in simulation output sufficiently close to the observed data. The accepted parameters are used to compute the approximate posterior distribution, ℙ(***θ***|𝓓^*obs*^). In order to assess how close the simulation and observed data are, we consider a range of summary statistics (detailed in Section 2.3.2) and an appropriate distance function (described in Section 2.3.3).

In order to quantify the performance of different summary statistics in inferring model parameters, we choose a uniform (uninformative) prior,

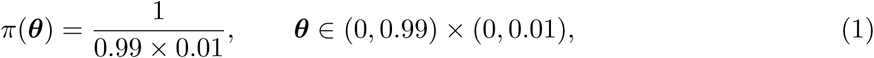

to ensure *P_m_* + *P_p_* ≤ 1.

#### 2.3.1. ABC rejection

Algorithm 1 describes the ABC rejection algorithm we use in this work. In Algorithm 1 we avoid directly specifying an acceptance threshold, *ϵ*, by accepting the 1^st^ percentile of the samples (ranked in terms of the distance between the simulated and observed data) and taking the number of samples, *K*, sufficiently large to obtain an accurate approximation to the true posterior. The accepted samples are used to estimate a posterior distribution using bivariate kernel density estimation with a Gaussian kernel [7].

##### Algorithm 1

ABC rejection algorithm.

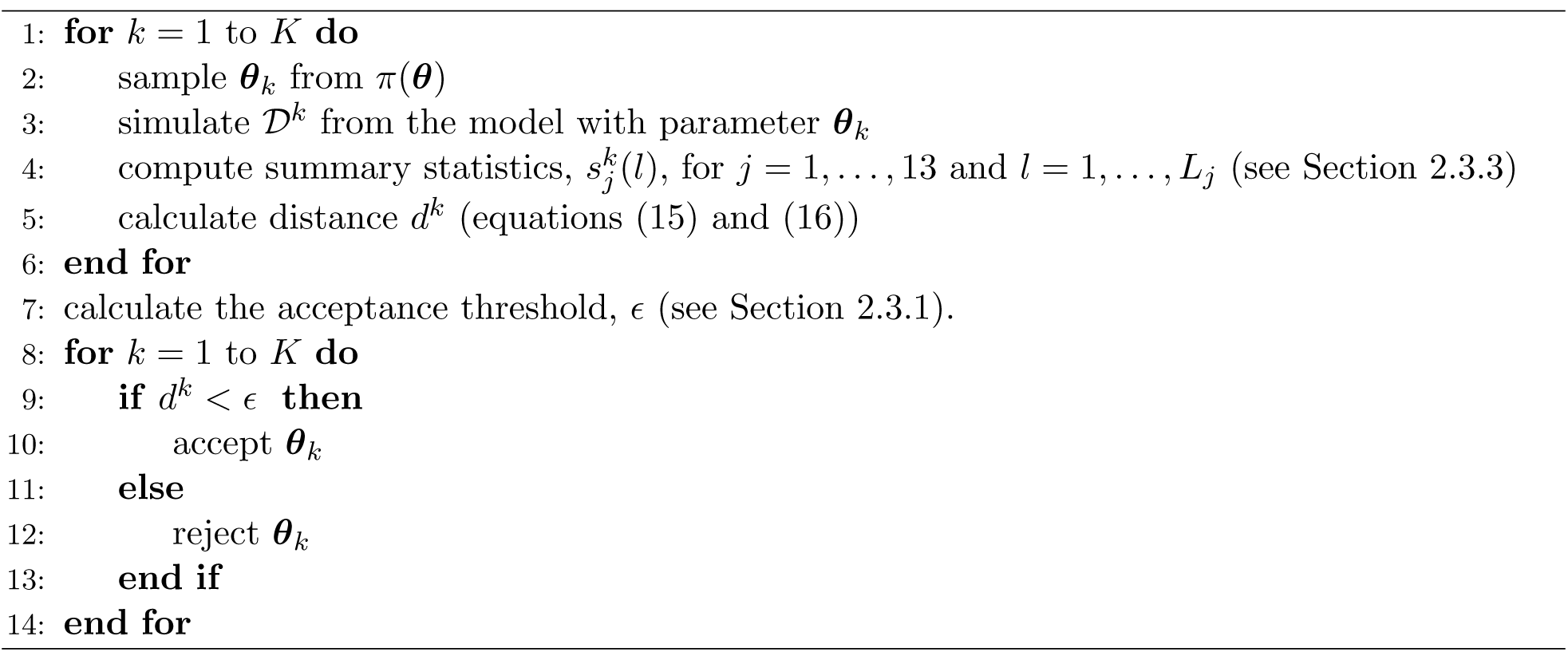

#### 2.3.2. Summary statistics

In order to compare observed and simulated data, we reduce the dimension of the data using summary statistics. How best to choose summary statistics for parameter inference is an ongoing research question, and some automated procedures have been developed for this choice, typically either through minimising a loss of information function [1], or maximising the gain of information [3], through a measure such as the Kullback-Leibler divergence [14].

*Kulback*-*Leibler divergence*. The Kullback-Leibler divergence is a measure of the relative difference in two continuous distributions, *F* and *G*, and defined as

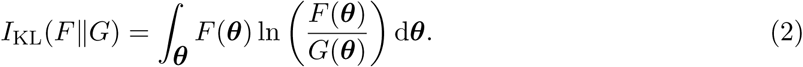

In this work, we generate bivariate posteriors and priors which are each functions of ***θ*** = (*P_m_*, *P_p_*). We discretise these distributions onto a fine mesh, 512 × 512 in size, for plotting. We compute the Kullback-Leibler divergence, *I_KL_*, by numerically integrating over both dimensions.

The observed data consists of the positions of all cells at the terminal time of the assay together with the tracks of five randomly chosen cells, *i* = 1,…, 5. We let *N*(*t_n_*) be the number of cells at time *t_n_* = *τ* × *n*, i.e. at the *n*^th^ iteration of a simulation, so that *n* = 288 corresponds to the final simulation time, *T* = 12hrs. We also let ***X**_i_*(*t_n_*) = (*X_i_*(*t_n_*),*Y_i_*(*t_n_*)) be the (*x*,*y*) lattice coordinates of cell *i* at time *t_n_*, *δX_ij_* (*t_n_*) = *X_i_*(*t_n_*) – *X_j_* (*t_n_*), and similarly for *δY_ij_*. We summarise these data using statistics motivated by random walk models [3, 12, 25], considering thirteen statistics in total, labelled as in Table 1.

**Table 1:** A list of the summary statistics and the corresponding notation adopted. Note, apart from cell trajectory statistics, all summary statistics are evaluated at the final time, *T*.

| Summary statistic | Notation |
| --- | --- |
| Number of cells | $N$ |
| Size of largest cluster, $k$-connected | $\kappa_k, k = 4, 8$ |
| Binning variance, bin size = $k$ | $Q_k, k = 2, 4, 8$ |
| Manhattan displacement of cell $i$ | $\ x\ $ |
| Manhattan tortuosity of cell $i$ | $\Gamma$ |
| Smallest gyration tensor eigenvalue | $\lambda$ |
| Correlation function | $C_{XY}(l)$ |
| Normalised two-dimensional correlation function | $\hat{C}_{XY}(l)$ |
| One-dimensional correlation function | $C_Y(l)$ |
| Normalised one-dimensional correlation function | $\hat{C}_Y(l)$ |

- The first summary statistic we consider is the final cell number, *N*(*T*).
- In order to assess whether cells are forming clusters, the second and third summary statistics we consider relate to the size of the largest cluster of cells, computed using the MATLAB function `bwconncomp` [19]. We consider either 4-connected clusters (cells orthogonally adjacent are part of a single cluster), *κ*_4_(*T*), or 8-connected clusters (cells diagonally adjacent are also part of a single cluster), *κ*_8_(*T*).
- Summary statistics four to six correspond to the binning variance [25], which quantifies the deviation from the average number of cells expected in quadrats of the domain, and is computed as

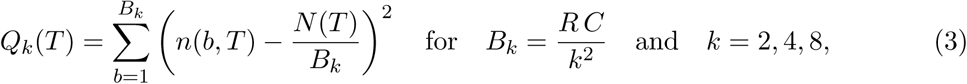

where *n*(*b*, *T*) is the number of cells in bin *b* at time *T*, and *B_k_* the number of bins when the bin width is *k*.
- As summary statistics seven and eight we consider trajectory statistics from the five tracked cells (labelled *i* = 1,…, 5): either the total Manhattan displacement (the sum of horizontal and vertical distances moved) of the cells, ∥*x*∥, where

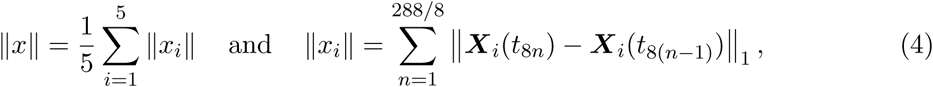

or the tortuosity of the trajectory, Γ, where

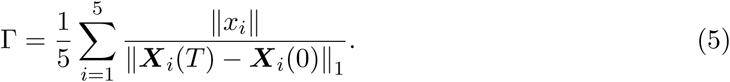
- Summary statistic nine is the smallest eigenvalue, λ, of the gyration tensor [23],

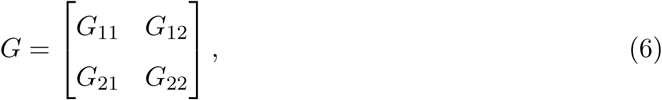

where

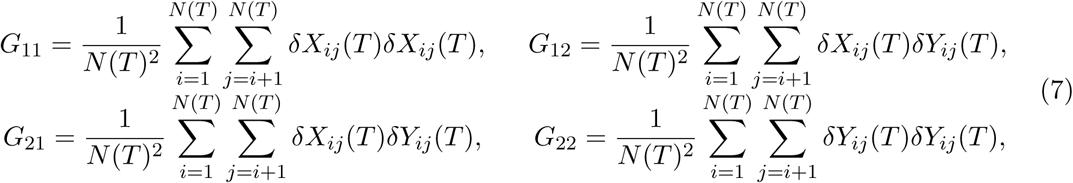

and the smallest eigenvalue quantifies the spread of cells along the lesser of the two principal axes.
- Summary statistics ten and eleven consider the distribution of pairs of cells, using the pairwise correlation functions, where the argument *l* indicates the number of pairs of cells separated by distance *l*. We consider pairwise correlations that measure only the vertical separation between cells, *C_Y_* (due to the heterogeneity of initial condition in the *y* direction), or the total separation of cells, *C_XY_*, where

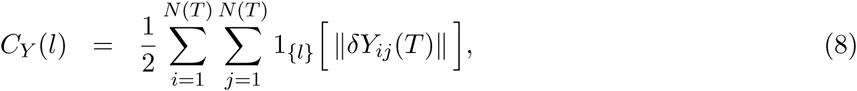

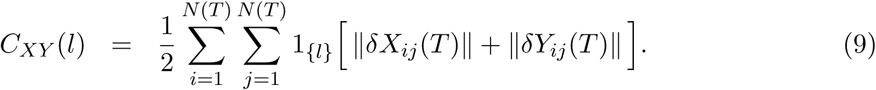
- Finally, we consider the correlation functions normalised by the expected number of pairs at each distance *l* = 1,…, *L*, which accounts for the density of cells in the domain,

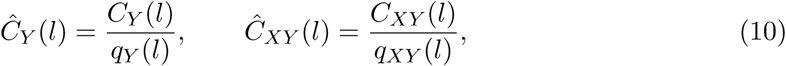

where

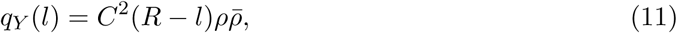

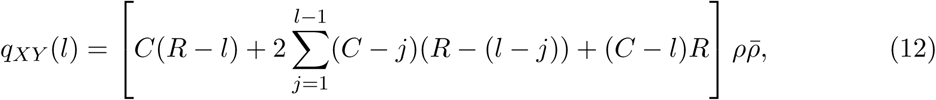

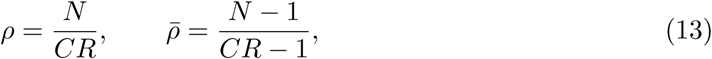

and *C* and *R* are the numbers of columns and rows in the lattice, respectively.

#### 2.3.3. Computing distances

Experiments are typically performed multiple times in similar conditions, so we simulate multiple data sets, and call each set a replicate. The observed data, 𝓓^*obs*^, consists of *M* replicates, 𝓓^*obs*^ = {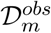: *m* = 1,…, *M*}. The simulated data, 𝓓, consists of *M* replicates and *K* samples, 𝓓 = {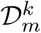: *m* = 1,…, *M*; *k* = 1,…, *K*}. From the simulated data we compute each summary statistic, *S_j_*(*l*), where *j* = 1,…, 13 denotes the thirteen different summary statistics, and *l* = 1,…, *L_j_* is the *l*^th^ element of statistic (*L_j_* = 24 for the pairwise correlation function statistics because there are 24 rows in the lattice, and *L_j_* = 1 otherwise).

First we average over the replicates,

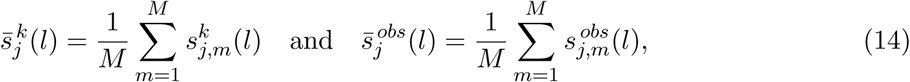

then we compute the median absolute deviation (MAD) for each statistic, *σ_j_* (*l*). This is defined for a univariate data set, *X* = {*X_i_*}, as MAD = median(|*X_i_* – median(*X*)|). We then compute the distances for each statistic as

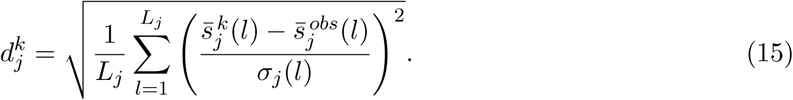

Note that this choice of distance function weights vectors of summary statistics of different lengths equally. This is of particular importance when performing parameter inference utilising combinations of summary statistics, as in this work. We consider combinations of *J* = 1, 2 or 3 summary statistics from the thirteen statistics listed in Table 1 to give

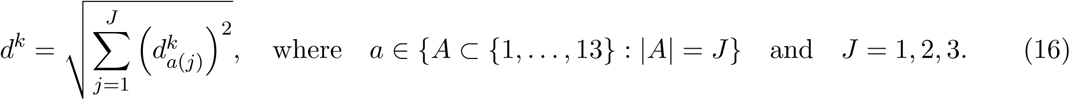

#### 2.3.4. Regression adjustment

After performing ABC rejection (and during ABC-DC) we perform regression adjustment to derive a more accurate estimate of the posterior distribution [4, 20, 21]. We assume that ***θ*** satisfies the following regression model,

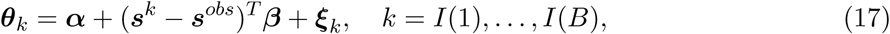

where *I* = {*k*: *d^k^* < *δ*}, *B* = |*I*|, ***s*** denotes the vector of summary statistics considered in the distance function (16) and the ***ξ**_k_* are uncorrelated gaussian random variables with zero mean and common variance ***σ***^2^. The regression model is solved to find the least-squares estimates, (***α̂***, ***β̂***), and the accepted parameter sets are adjusted according to

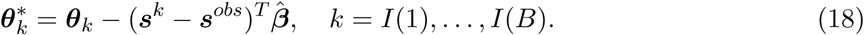

#### 2.3.5. Data-cloning ABC

Data-cloning ABC involves considering a data-set, 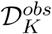,containing *K* clones of the experimental data, 𝓓^*obs*^, that is, 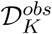 = (𝓓^*obs*^, 𝓓^*obs*^,…, 𝓓^*obs*^), where each clone is assumed independent of the others. The likelihood of the cloned data is then [15, 21],

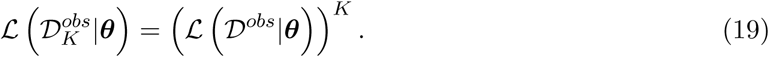

Hence the posterior distribution resulting from cloned data satisfies

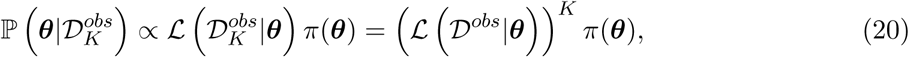

and the MLE of ***θ*** is equivalent to the mean of ℙ(***θ***|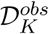) as *K* → ∞ [15]. Intuitively, we can see this by reasoning that, if all of the independent model-generated data-sets are close to the experimental data, we are *K* times as likely to have selected a sensible candidate parameter.

The ABC-DC algorithm was proposed by Picchini *et al*. [21], and it uses ABC Markov chain Monte Carlo [17]. The algorithm works in two stages, first the acceptance threshold, *ϵ*, is decreased according to a tolerance scheme {*ϵ*_1_, *ϵ*_2_,…, *ϵ_P_*}, and then the number of clones, *K*, is increased according to a clones scheme (*K*_*P*+1_,*K*_*P*+2_,…, *K_P+Q_*}, where subscripts index the population number. The MLE is approximated by averaging the final parameter population, and the 90% credible interval can be used for comparison to ABC rejection.

For each sample, a proposal ***θ***^#^ is accepted or rejected according to the acceptance probability

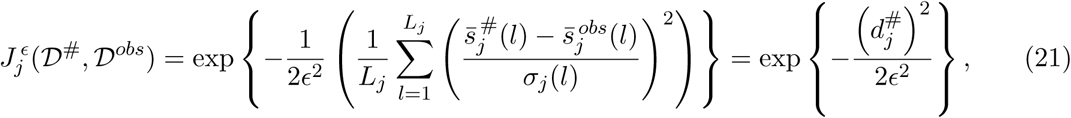

which approximates a Gaussian centred on the experimental observation with variance *ϵ*, weighted according to the MAD, *σ_j_*(*l*), of the summary statistic, *j* under consideration, where the MAD is estimated using ABC rejection.

The ABC-DC method is described in full in Algorithm 2. The number of samples, *r*, for the decreasing tolerance stage is denoted by (*r*_1_,…. *r_P_*}, and (*r*_*P*+1_,…,*r_P+Q_*} denotes the number of samples for the increasing clones stage. The scheme we use has *P* = 5 and *Q* = 4, and other variables are as follows:

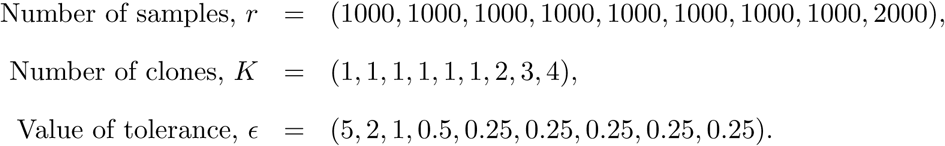

## 3. Results

Our aim in this work is to provide insights into how both the choices of experimental geometry and summary statistics impact the quality of the posteriors generated using ABC for experiments that are typically used to characterise growing and spreading cell populations. We generate *in silico* (observed) data using the model outlined in Section 2.1 and Section 2.2 for the range of experimental designs illustrated in Figure 1(c), and quantify the quality of the posteriors resulting from the use of ABC rejection with various summary statistics using the Kullback-Leibler divergence between the prior and posterior distributions (details of the methods used can be found in Section 2.3). In Section 3.1 and Section 3.2 we ask whether it is possible to quantify, using only a single summary statistic of the data, the relative contributions of proliferation and motility in a number of cell spreading experiments. In Section 3.3 we explore the extent to which parameter estimates can be improved using multiple summary statistics. Finally, in Section 3.4 we demonstrate the use of ABC-DC to efficiently find MLEs, and compare these estimates to those generated using ABC rejection.

### Algorithm 2

ABC-DC algorithm.

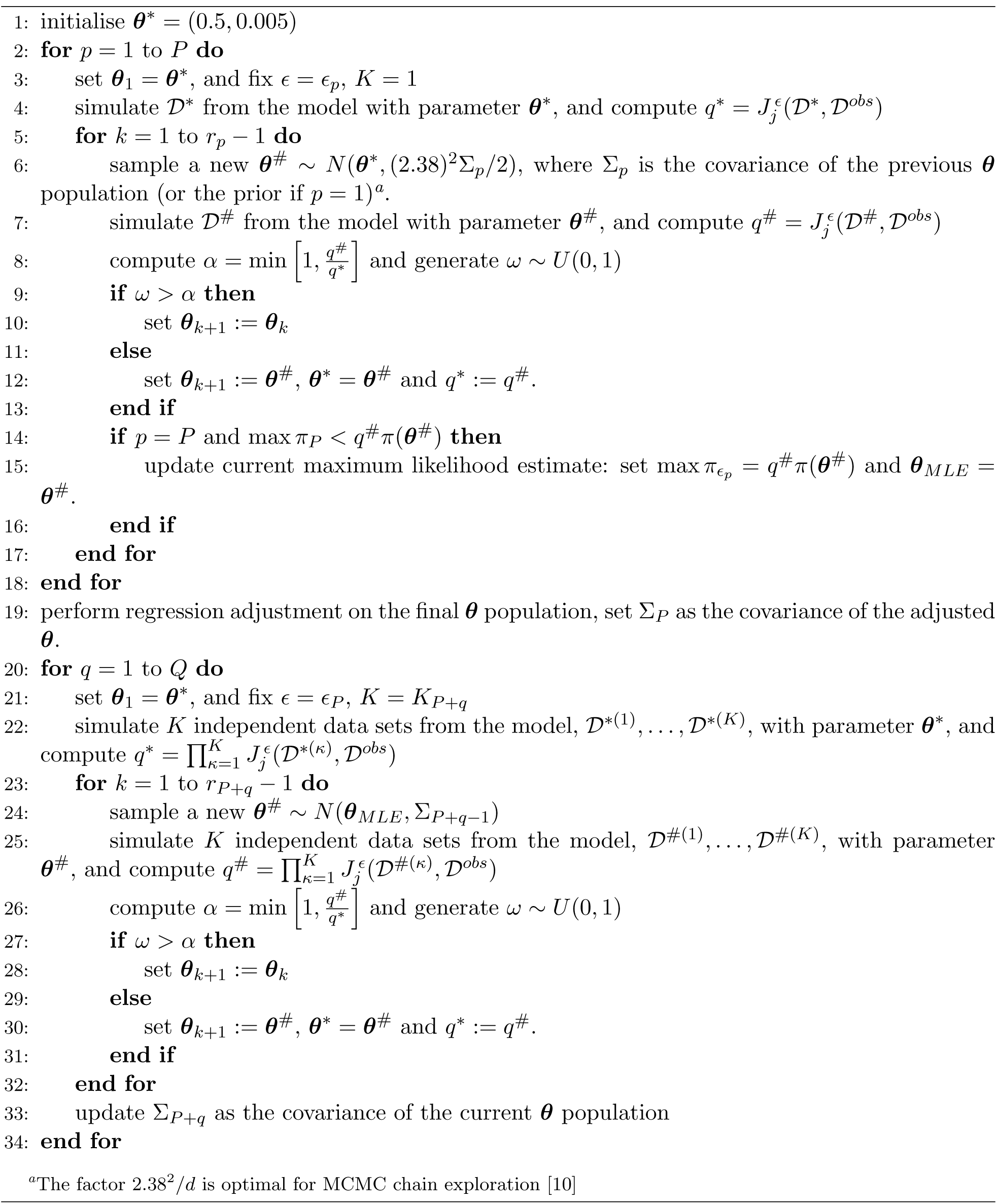

### 3.1. Can we infer both model parameters using a single summary statistic?

We first ask whether we can jointly infer both model parameters, *P_m_* and *P_p_*, using a single summary statistic in the distance function (16). To do this, we perform ABC rejection on *in silico* (observed) data gathered from simulations with *P_m_* = 0.25 and *P_p_* = 0.0025, where we replicate different experimental designs by varying the model initial conditions: we initialise the model with 24 cells either in all 24 rows, in 12 rows, or in six rows, of the domain, as shown in Figure 1(c). Note that this means we change the average density within the region into which cells are initialised, but that the overall cell number, and hence the average total density, is constant. Figure 2(a)-(f) shows the posteriors generated for selected summary statistics of the data. The average information gain, and its deviation, for each of the three experimental designs and all summary statistics, is summarised in Figure 2(g).

**Figure 2:**
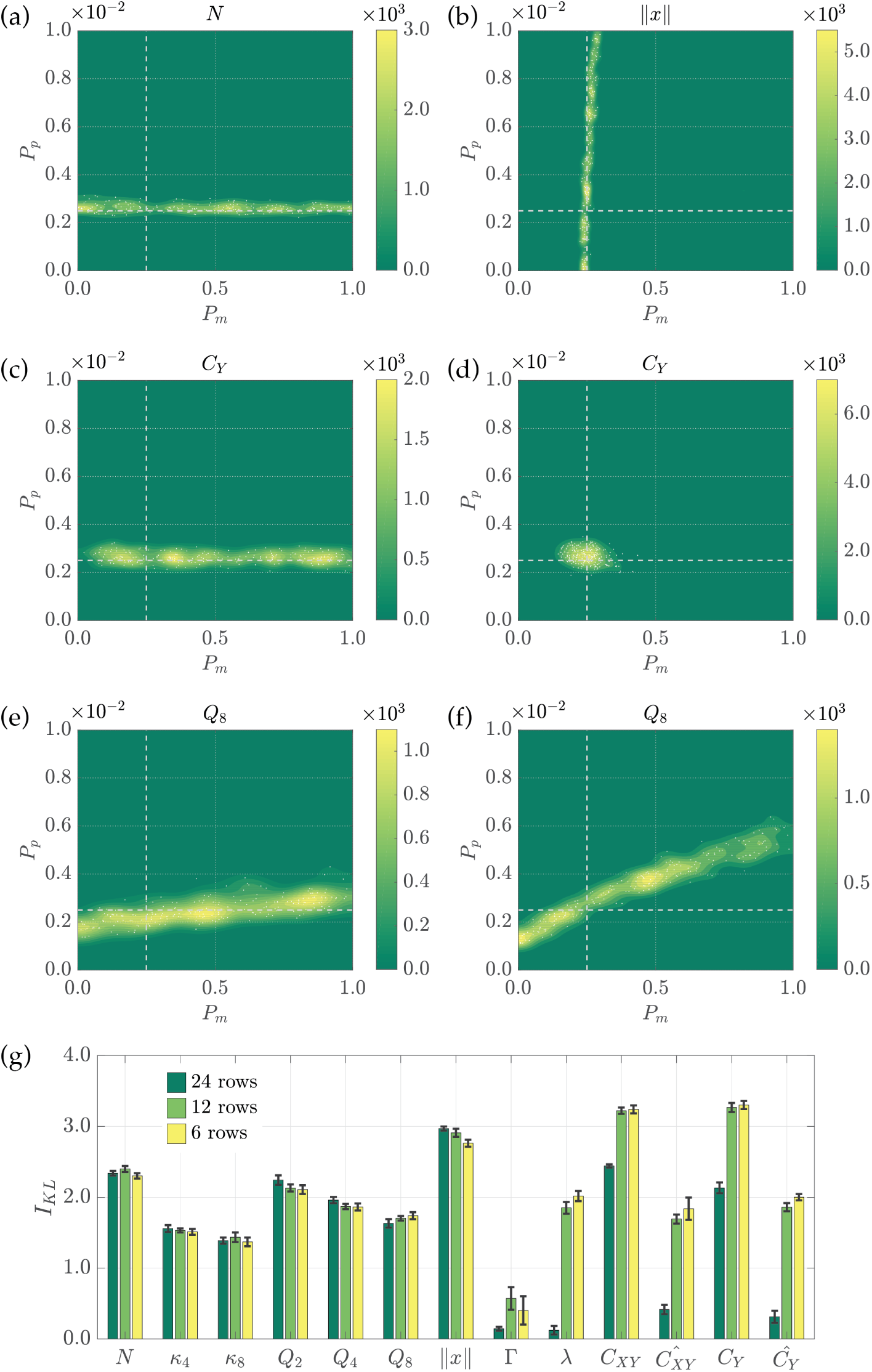
Posteriors generated using a single summary statistic as the experimental design is varied. The parameters used to generated the *in silico* (observed) data are indicated using dashed white lines. (a),(b) The posterior distributions corresponding to the most informative summary statistic for inferring *P_p_* ((a), *N*) or *P_m_* ((b), ∥*x*∥) for the growth-to-confluence assay (cells initialised uniformly at random over 24 rows). (c),(d) The posterior distributions generated using the summary statistic *C_Y_* for the growth-to-confluence assay (c) and the scratch assay (d) (cells initialised uniformly at random over six rows). (e),(f) The posterior distributions for summary statistic *Q*_8_ for the growth-to-confluence assay (e), and the scratch assay, (f). (g) The information gain in moving from the prior to the posterior for each summary statistic and for each experimental design, where the error bars denote the standard deviation.

Figure 2(a)-(b) shows that for a growth-to-confluence assay design (where cells are initialised across all 24 rows of the domain), an accurate estimate of *P_p_* can be obtained using the number of cells, *N*, as a summary statistic, and an accurate estimate of *P_m_* can be obtained using the Manhattan displacement, ∥*x*∥, as a summary statistic. However, no single summary statistic can be used to infer both *P_p_* and *P_m_* with any degree of confidence. Figure 2(c)-(d), however, demonstrates that, in contrast to a growth-to-confluence assay design, using a scratch assay design (where cells are initialised only in six rows of the domain) enables both *P_m_* and *P_p_* to be accurately estimated using a single summary statistic, for example the one-dimensional correlation function summary statistic, *C_Y_*. Finally, Figure 2(e)-(f) demonstrates how the predicted posterior can vary for a single summary statistic as the experimental design is changed. We use as an example the binning variance statistic, *Q*_8_, which is the deviation in cell numbers from the mean when the domain is divided into 8×8 bins. Increasing the rate of movement, *P_m_*, means clusters are broken up more rapidly, whilst increasing the rate of proliferation, *P_p_*, makes clusters larger. As a result, *in silico* (observed) data generated using small *P_m_* and *P_p_* results in the same binning variance as data generated with large *P_m_* and *P_p_*. The effect is more pronounced for a scratch assay design due to the initial confinement of cells.

Figure 2(g) summarises the quality of the predicted posteriors resulting from the use of each of the summary statistics listed in Table 1 for each experimental design considered in Figure 1(c). Our results demonstrate that statistics based on cell numbers provide the same or less information as cells are increasingly confined initially, whereas the opposite is true for the summary statistics that relate to correlations in cell positions. Both model parameters, *P_m_* and *P_p_*, can be inferred using a single summary statistic, without the need for cell trajectory information, but only when cells are initially confined to a scratch-assay-like geometry. For growth-to-confluence assays, cell trajectory data is necessary to accurately infer both model parameters.

### 3.2. What is the impact of the initial number of cells on the posteriors?

Next we consider how changing the initial number of cells in the assay (24, 48 or 72 cells) affects the quality of the predicted posterior distribution. We consider either a growth-to-confluence assay (cells initialised uniformly at random across the domain, Figure 3(a)-(c)) or a scratch assay design (cells initialised uniformly into six rows of the domain, Figure 3(d)-(f)). Figure 3(a)-(c) demonstrates that, for the growth-to-confluence assay, increasing the initial cell number does not facilitate accurate inference of both model parameters using a single summary statistic. In line with the results presented in Figure 2, Figure 3(d)-(f) indicates that both model parameters can be accurately estimated using a single summary statistic when the scratch assay is used; however, increasing the initial cell number does not significantly increase the quality of the posterior. We quantify the quality of the estimated posterior for each summary statistic and each experimental design in the bar plots in Figure 3(c),(f). For almost every summary statistic the quality of the posterior increases as the initial cell number increases, likely due to a reduction in the variance of each summary statistic.

**Figure 3:**
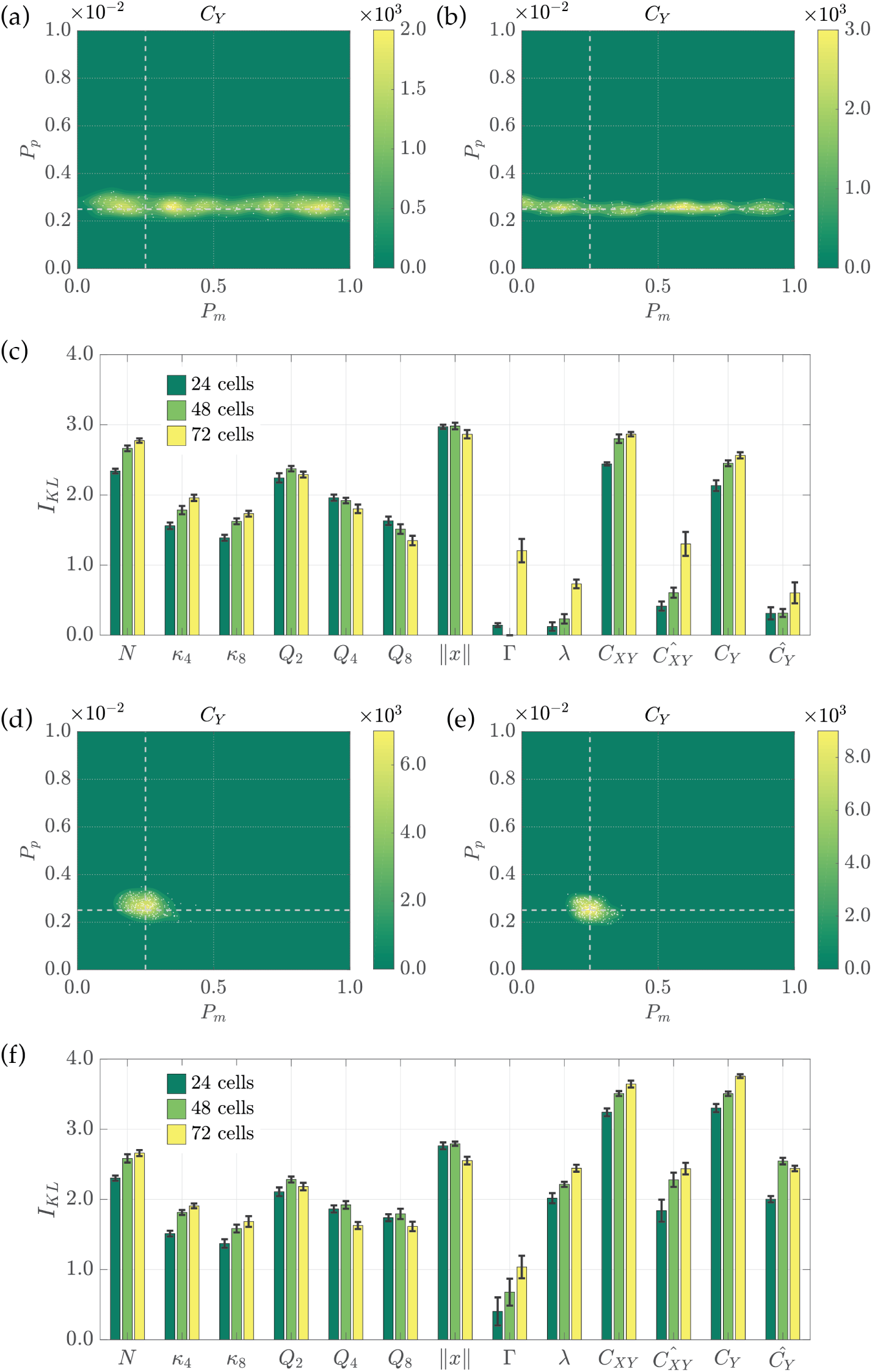
Posteriors generated using a single summary statistic as the initial cell number is varied. The parameters used to generated the *in silico* (observed) data are indicated using dashed white lines. (a)-(c) Results for a growth-to-confluence assay generated using 24 cells (a) and 72 cells. (d)-(f) Results for a scratch assay generated using 24 cells (d) and 72 cells (e). The posterior distributions in (a),(b) and (d),(e) are generated using the pairwise correlations summary statistic *C_Y_*. (c),(f) The information gained in moving from the prior to the posterior for each summary statistic for each experimental design, where the error bars denote the standard deviation.

### 3.3. Are trajectory statistics necessary for maximal information gain?

It is natural to ask whether multiple summary statistics can be combined to increase the quality of the estimated posteriors and, further, whether combining different summary statistics can avoid the need to collect cell trajectory data to accurately infer parameters of a growth-to-confluence assay. To answer this question, we considered all combinations of two or three summary statistics from the 13 under consideration, computing the resulting posteriors and information gain for each combination. Figure 4 demonstrates that, for a growth-to-confluence assay, cell trajectory data are necessary to generate a posterior that allows for accurate estimation of both parameters *P_m_* and *P_p_*(Figure 4(a),(b)). In particular, trajectory information is necessary to estimate the motility parameter, *P_m_*. It is interesting to note that, in both cases (with and without trajectory information), the number of cells summary statistic, *N*, is included in the best-performing two-summary-statistic combinations, likely because it facilitates accurate estimation of the proliferation parameter, *P_p_*. For a scratch assay, cell trajectory information is not necessary to accurately infer both model parameters, however, as expected, including cell trajectory data enables more accurate inference of the cell motility parameter, *P_m_*. Figure 4(e) quantities the quality of the estimated posteriors for a range of experimental designs, initial cell densities and for combinations of one, two or three summary statistics. Results demonstrate that little extra information is gained by considering more than two summary statistics and that, as expected, accurate parameter inference for growth-to-confluence assays requires cell trajectory data.

**Figure 4:**
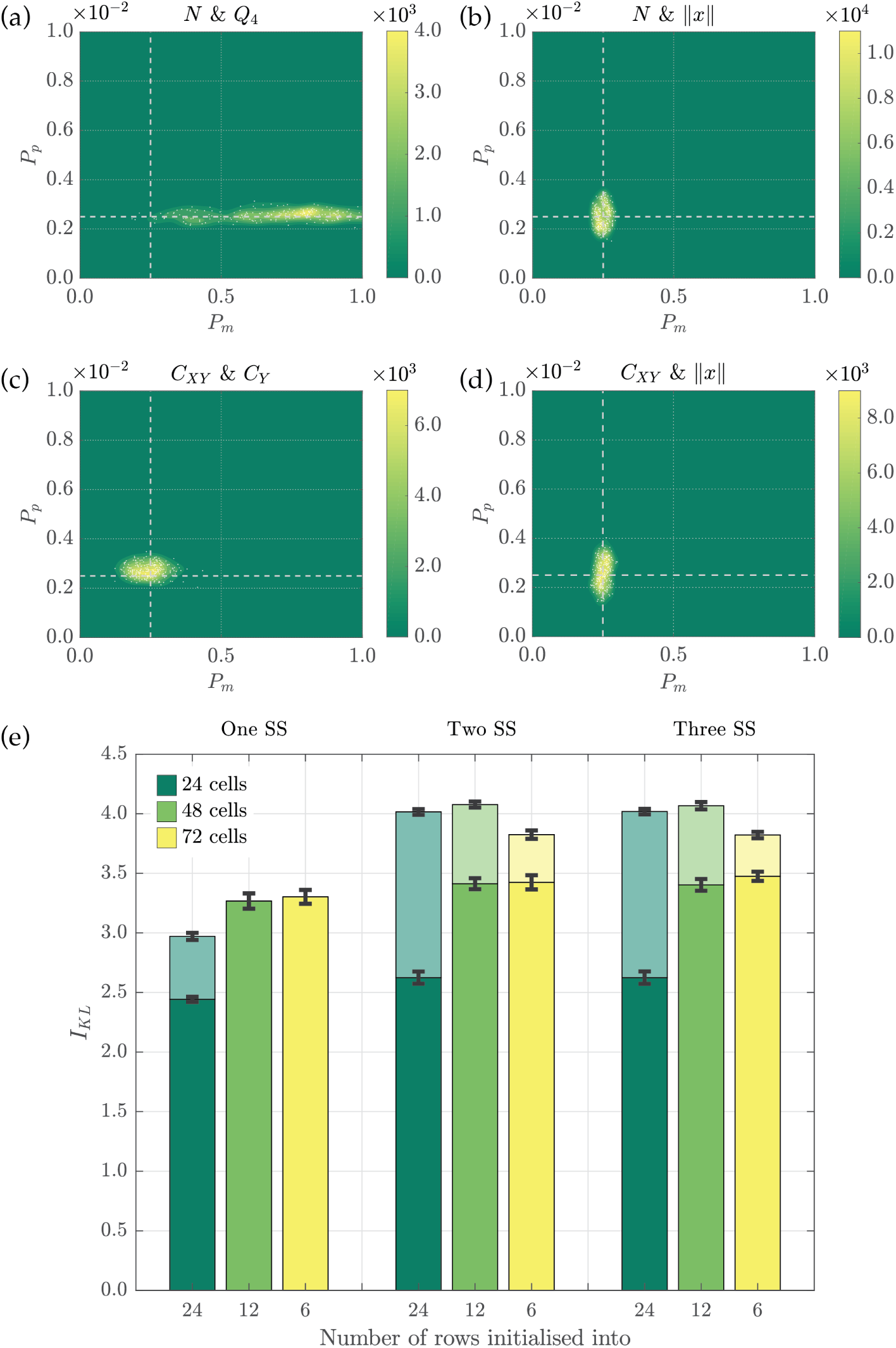
Posterior distributions and information gain when considering combinations of summary statistics (SS). (a)-(d) show posteriors resulting from the most informative two-summary-statistic combination when considering either a growth-to-confluence assay (a),(b), or a scratch assay, (c),(d), excluding trajectory information (a),(c), or including trajectory information (b),(d). (e) The information gain in moving from the prior to the posterior for each of the best performing summary statistic combinations, varying the experimental design, and excluding trajectory information (dark shading) or including trajectory information (light shading), where the error bars denote the standard deviation.

### 3.4. Can data-cloning ABC provides an efficient means to estimate model parameters?

Finally, we investigate whether the ABC-DC algorithm proposed by Picchini *et al*. [21] can provide an efficient means to estimate model parameters. ABC-DC essentially involves using a number of copies of the observed data (called “clones”) that are assumed to be independently generated data sets. ABC-DC uses ABC Markov chain Monte Carlo [17] over two stages: first the acceptance threshold, *ϵ*, is decreased; and then the number of clones, *K*, is increased. This has the impact of concentrating the posterior around the MLE, with the posterior mean an approximation to the MLE, and the 90% credible interval available for quantification of the uncertainty in parameter predictions. Figure 5 shows the results of using ABC-DC with the two-dimensional pairwise correlation function, *C_XY_*, to estimate posterior distributions for both *P_m_* and *P_p_*. Figure 5(a),(c) shows how the posterior changes as the number of clones is increased, relative to ABC rejection, whilst Figure 5(b),(d) shows the Markov chains of both *P_m_* and *P_p_*. Our results indicate that, as expected, the ABC-DC algorithm results in a tighter posterior focussed approximately about the MLEs found using ABC-DC with a single instance of the data, *K* = 1. To understand how the MLEs generated using ABC-DC differ from those generated using ABC rejection we look at the posteriors generated using each algorithm together with three summary statistics: the number of cells, *N*; the two-dimensional pairwise correlation statistic, *C_xy_*; and the Manhattan displacement, ∥*x*∥. We calculate MLEs and the 5% and 95% empirical percentiles of the accepted parameters. Results in Table 2 indicate that the MLEs are similar for both ABC rejection and ABC-DC, though the credible intervals for ABC-DC are much smaller, and not a good representation of the true uncertainty in each parameter. However, ABC-DC can provide significant computational savings in estimation of the MLE, with maximal efficiency displayed with summary statistics that are relatively computationally demanding to compute, such as *C_XY_*.

**Figure 5:**
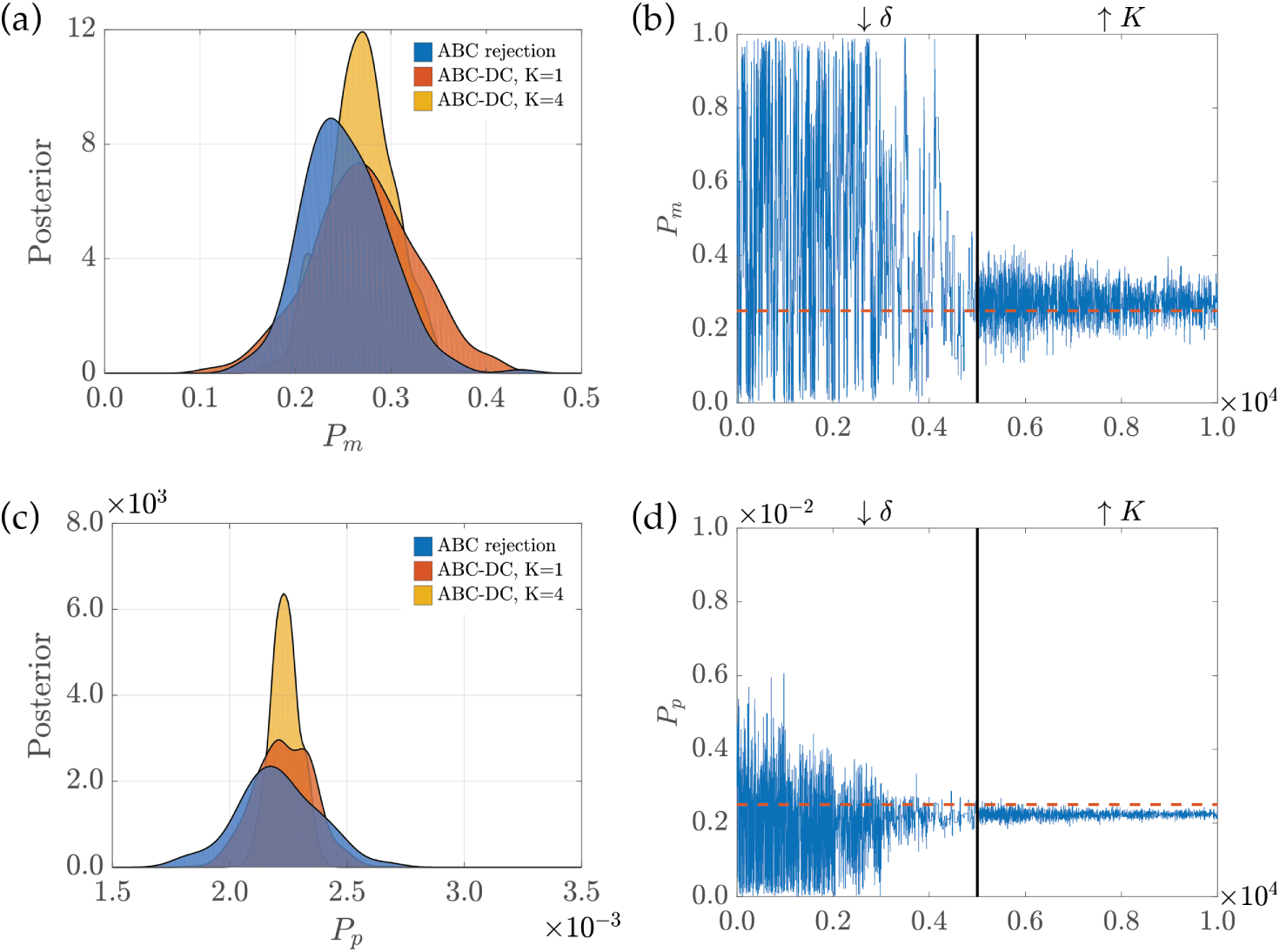
Example results from the ABC-DC algorithm for a scratch assay using the two-dimensional pairwise correlations statistic, *C_XY_*. (a),(c) show estimates of the posteriors for *P_m_* and *P_p_* generated using kernel density estimation. We compare the posteriors for *P_m_* and *P_p_* generated using ABC-DC (yellow), ABC-DC (before the clones numbers are increased) (red) and ABC rejection (blue). (b),(d) show the trajectories of parameters accepted during the Markov chain Monte Carlo step.

**Table 2:** ABC-DC for a scratch assay experiment. The MLEs and the 5%-95% empirical percentiles are shown for parameters accepted using ABC rejection and ABC-DC, for different summary statistics. The speed up of the ABC-DC method over ABC rejection is also given.

| Statistic: Parameter | ABC rejection MLE (90% CI) | ABC-DC MLE (90% CI) | Speed up |
| --- | --- | --- | --- |
| $\ x\ : P_m$ | 0.250 (0.231, 0.300) | 0.247 (0.239, 0.255) | 1.65 |
| $C_{XY}: P_m$ | 0.239 (0.174, 0.310) | 0.236 (0.186, 0.280) | 4.50 |
| $C_{XY}: P_p(\times 10^{-2})$ | 0.241 (0.215, 0.270) | 0.228 (0.219, 0.236) | 4.50 |
| $N: P_p(\times 10^{-2})$ | 0.243 (0.220, 0.282) | 0.246 (0.235, 0.257) | 1.72 |

## 4. Discussion

Mechanistic models together with experimental data from cell biology assays provide an excellent opportunity to characterise the extent to which cell proliferation and cell motility can drive the growth and expansion of a cell colony. To be able to draw quantitative distinctions between the relative contributions of cell proliferation and motility for different cell types and biological scenarios requires the accurate inference of model parameters using quantitative summaries of experimental data. In this work we have applied ABC to explore the extent to which different experimental designs and quantitative summary statistics impact our ability to accurately infer parameters of a simple mechanistic model of cell growth and spreading. Our results suggest that, for growth-to-confluence experiments, cell trajectory information, in addition to summary statistics such as cell numbers and spatial correlations, are required for accurate parameter inference. On the other hand, for a scratch assay geometry, it is possible to accurately infer both motility and proliferation parameters without recourse to tracking individual cells over multiple frames of a microscopy video. Since cell tracking provides the bottleneck in the analysis of many experimental studies, this renders experimental geometry an important aspect to consider the design of experiments.

In addition, our results demonstrate that increases in the initial cell number in cell biology assays can increase the quality of estimated posterior distributions by decreasing noise in the summary statistics, a result also found for a similar model by Ross and co-workers [22]. We also demonstrated that posteriors of significantly increased quality can be generated using a combination of two summary statistics, but that further increases in the number of summary statistics provided little increase in the quality of the posteriors. Finally, we have shown that the ABC-DC approach provides an efficient means to approximate the MLEs of model parameters, with significant improvements in computational efficiency provided where the summary statistics are time-consuming to calculate.

### 4.1. Predictions for experimental data

We can use our model to infer parameter distributions for different cell types in a growth-to-confluence assay, and subsequently make predictions about long-time behaviours, such as the time taken to reach confluence. In Figure 6 we use data from the growth-to-confluence experiments detailed in [25] to infer motility and proliferation rates for two different cell types: 3T3 fibroblast cells and MDA MB 231 breast cancer cells. Figure 6(a),(b) shows the posteriors generated using ABC and the cell number, *N*, and Manhattan displacement, ∥*x*∥, as summary statistics, and the priors

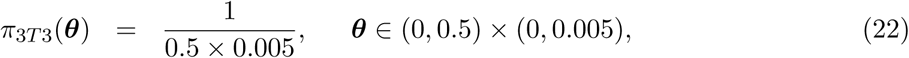

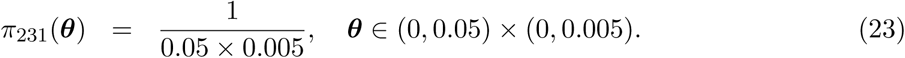

**Figure 6:**
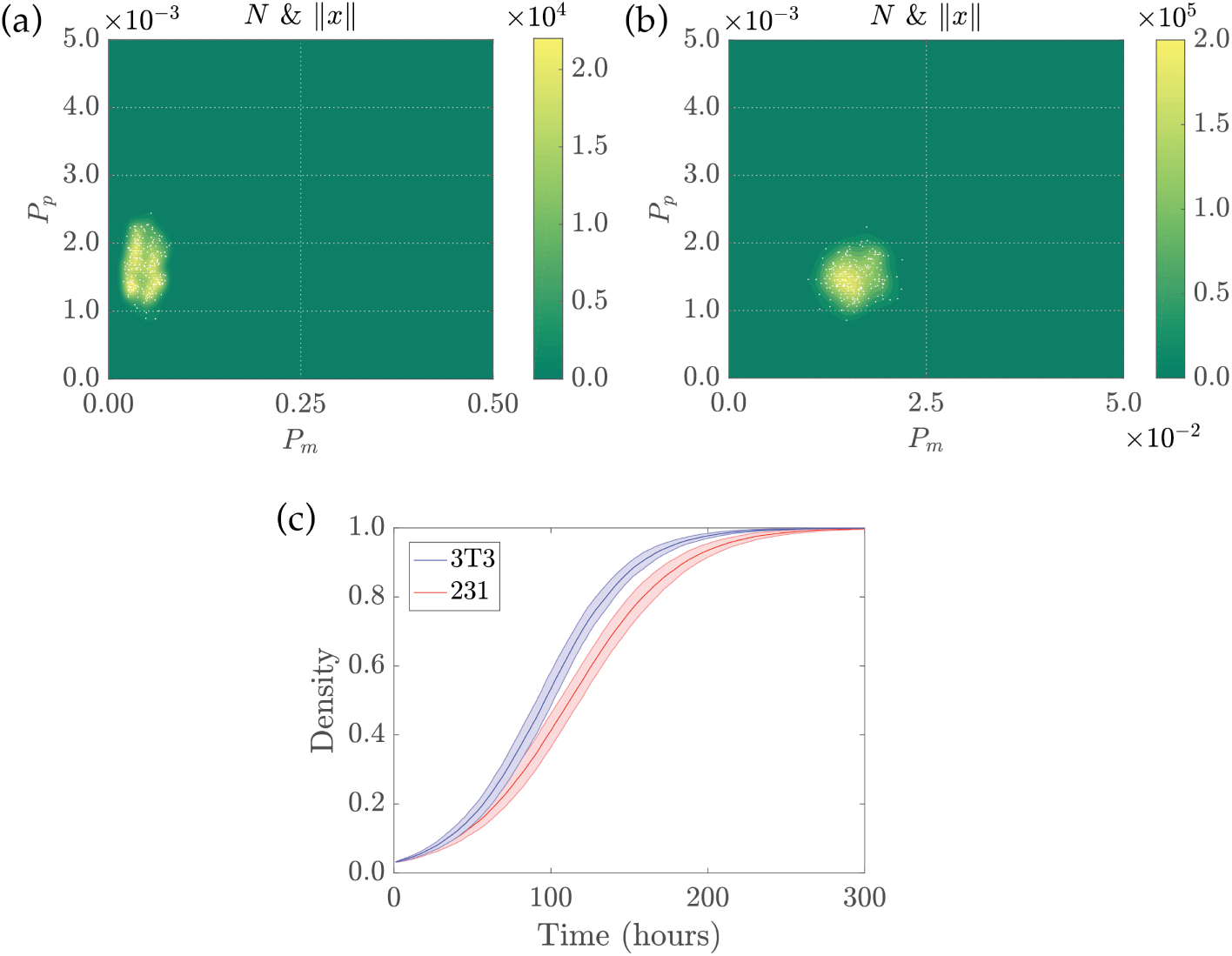
Posterior distributions and predictions for long-term dynamics. (a),(b) Posterior distributions for 3T3 fibroblast and 231 breast cancer cell lines, respectively. (c) Long-time predictions of the cell density generated using the mechanistic model described in Section 2.1, with 24 cells initially distributed uniformly at random within the domain and parameters as detailed in Section 4.1. The shaded areas represent the mean behaviour plus or minus one standard deviation.

For the 3T3 fibroblast cell line we obtain *P_m_* = 0.0464 ± 0.0148 and *P_p_* = 0.00165 ± 0.000325, whilst for the MDA MB 231 breast cancer cells we obtain *P_m_* = 0.0159 ± 0.00233 and *P_p_* = 0.00150 ± 0.000249. In each case we give the posterior mean and the standard error. In real terms, for the 3T3 cells this gives a diffusivity of *D* = 97.875 ± 31.219 *μ*m^2^ hr^−1^ and a proliferation rate of λ = 0.0396 ± 0.0078 hr^−1^, whilst for the 231 cells we have *D* = 33.539 ± 4.915 *μ*m^2^ hr^−1^ and λ = 0.0360 ± 0.0059 hr^−1^. All estimated parameters are consistent with predictions provided in [25]. In Figure 6(c),(d) we show long-time predictions for both cell populations. Although the proliferation rates are with 10% of each other, the disparity in diffusivities means that the 3T3 cells reach confluence much quicker. For example, the model predicts that the 3T3 cells fill 99% of the lattice after approximately 205 hours whilst the 231 cells take on the order of 244 hours, approximately 30% longer.

### 4.2. Conclusions

In summary, our results suggest that experimental design choices that incorporate initial spatial heterogeneities in cell positions facilitate parameter inference without the requirement of cell tracking, whilst those that seed cells uniformly initially require cell tracking for accurate parameter inference. As cell tracking is often a major technical limitation many experimental studies of this type, our recommendations for experimental design choice could lead to significant potential time and cost savings in the analysis of these kinds of commonly-used experiments.

#### Data accessibility

Code used to generate the data are available from https://github.com/andrew-parker/Impact-of-experimental-design.

## Author contributions

AP, MJS and REB conceived the study, which was performed by AP. AP, MJS and REB wrote the manuscript.

## Acknowledgements.

AP would like to thank the UK’s Engineering and Physical Sciences Research Council (EPSRC, EP/G03706X/1) for funding through a studentship at the Systems Biology programme of The University of Oxford’s Doctoral Training Centre. MJS appreciates support from the Australian Research Council (DP170100474). REB is a Royal Society Wolfson Research Merit Award holder and would like to thank the Leverhulme Trust for a Research Fellowship.

